# Guam amyotrophic lateral sclerosis/parkinsonism-dementia complex (ALS/PDC) features CTE-like tau seeds in brain and spinal cord

**DOI:** 10.64898/2025.12.22.696002

**Authors:** Nil Saez-Calveras, Bert M. Verheijen, Nabil Morgan, Elizabeth Hill, Prarthna Chabria, Skyler Taylor, Kiyomitsu Oyanagi, Akiyoshi Kakita, Yuyu Song, Lukasz A. Joachimiak, Jaime Vaquer-Alicea, Marc I. Diamond, Ying Lu

**Affiliations:** Center for Alzheimer’s and Neurodegenerative Diseases, Peter O’Donnell Jr. Brain Institute, University of Texas Southwestern Medical Center, Dallas, TX, USA; Department of Neurology, University of Texas Southwestern Medical Center, Dallas, TX, USA; Department of Systems Biology, Harvard Medical School, Boston, MA, USA; Department of Neurology, Massachusetts General Hospital, Boston, MA, USA; Division of Neuropathology, Department of Brain Disease Research, Shinshu University School of Medicine, Matsumoto, Nagano, Japan; Department of Pathology, Brain Research Institute, Niigata University, Niigata, Japan

**Keywords:** amyotrophic lateral sclerosis-parkinsonism dementia complex (ALS/PDC), neurodegenerative disease, tau, seeding, alanine scan

## Abstract

Amyotrophic lateral sclerosis/parkinsonism-dementia complex (ALS/PDC) is a fatal neurodegenerative disorder that was once hyperendemic in the island of Guam (Mariana Islands, US) and a few other Pacific locales. Despite extensive investigations into its origins, the etiology of ALS/PDC remains unclear. ALS/PDC is, at the neuropathology level, characterized by tau-dominant multiple proteinopathy in brain and spinal cord. It was recently reported that Guam ALS/PDC brain extracts exhibit tau seeding activity in fluorescence resonance energy transfer (FRET)-based biosensor cells. To build upon those findings and explore the nature of tau seeds in ALS/PDC in more detail, we used an alanine mutational scanning (Ala scan) approach to determine the seeding profile of tau in nervous tissues of Guam ALS/PDC cases. First, we confirmed the detection of tau seeding activity in ALS/PDC brain samples in tau biosensor cells. Notably, we could also detect potent tau seeding activity in spinal cord. Subsequent Ala scan assays demonstrated that ALS/PDC tau displays an aggregate incorporation pattern that resembles that of chronic traumatic encephalopathy (CTE)-type tau. This result is consistent with recent electron cryo-microscopy studies of tau, which revealed that ALS/PDC tau filaments are predominantly of the CTE-type. The structural characteristics and seeding behavior of ALS/PDC tau, as well as the regional distribution of tau pathology at post-mortem, suggest that ALS/PDC is a CTE-like tauopathy.

**Significance Statement:** Neurodegenerative tauopathies are characterized by proteinaceous deposits containing microtubule-associated tau in nervous tissue. Emerging evidence suggests that disease-associated tau proteins adopt abnormal, self-propagating conformations characteristic of prions. Here, we employed alanine mutational scanning (Ala scan) to determine the nature of prion-like tau seeds in ALS/PDC, a mysterious disorder that occurred formerly in high incidence in certain regions in the western Pacific. We show that the Ala scan incorporation profile of ALS/PDC tau is similar to that of abnormal tau proteins in chronic traumatic encephalopathy (CTE). The findings lend support to the idea that ALS/PDC can be classified structurally as a CTE-like tauopathy. This work may have important implications for our understanding of ALS/PDC as well as common neurological disorders beyond the Pacific.

## Introduction

Amyotrophic lateral sclerosis/parkinsonism-dementia complex (ALS/PDC) is a rare neurodegenerative disorder of unknown cause that previously occurred in high incidence in the Pacific Island of Guam (Mariana Islands, US) (1–5) as well as other geographic foci in the western Pacific, i.e., the Kii peninsula of Japan (6, 7) and the southern coastal region of Papua (8, 9). The disease presents clinically as progressive motor neuron disease (ALS), parkinsonism with dementia (referred to as parkinsonism–dementia complex or PDC), or a combination thereof. Pacific ALS appears to be similar to ALS occurring elsewhere. PDC, however, seems to be a unique disorder found only in the western Pacific.

Research into ALS/PDC has been primarily stimulated by hypotheses as to cause. It was once hoped that understanding the cause of ALS/PDC in the Pacific isolates would provide insight into analogous neurological disease encountered throughout the world (e.g., idiopathic ALS). We will concentrate here on the Guam cluster of disease, i.e., the focus that harbors the most intensively studied ALS/PDC variant. Although various causes have been proposed for ALS/PDC in Guam, such as genetic factors, infections, and exposure to toxins, no conclusive evidence has been found for any of those (10). A remarkable decrease in disease incidence has been seen with increased westernization (11, 12) and migration studies have indicated that the disease could also develop after long-term residence in Guam (13). These observations hint at some role for exogenous factors in ALS/PDC etiopathogenesis.

At the neuropathology level, Guam ALS/PDC is characterized by widespread neurofibrillary tangles (NFTs) composed of hyperphosphorylated microtubule-associated tau protein (14–16), often in combination with other protein abnormalities, e.g., intracytoplasmic phospho-TAR DNA-binding protein of 43 kDa (pTDP-43) aggregates (17–19) (**Fig. S1**). Protein aggregates are found in brain and spinal cord tissues, in both neurons and glial cells in gray and white matter (20–25). NFTs in ALS/PDC brains are composed of all six tau isoforms (3R + 4R) and are prominent in superficial cortical layers, i.e., cortical layers II/III (26). Of note are the occurrence of granular/fuzzy astrocytes and subpial thorn-shaped astrocytes localized in the depth of cortical sulci (27). α-synucleinopathy can also be occasionally observed (28–30), but β-amyloid (Aβ) plaques are mostly absent and are not considered a defining feature of ALS/PDC [a summary of neuropathology can be found in reference (31)]. It has been reported that NFTs are also frequently present in nervous tissues of Guam non-neurological control subjects at post-mortem (32, 33), likely indicating a presymptomatic state. It is conceivable that shared genetic risk and/or exposure to a common etiologic agent resulted in a certain degree of background pathology in the Guam populace. ALS/PDC cases in the Kii cluster show neuropathological features that are largely comparable to those in Guam ALS/PDC (34). No necropsies were performed on ALS/PDC cases in the Papua cluster and it is therefore unknown whether those cases are neuropathologically similar to ALS/PDC in the other geographic isolates.

A growing body of evidence suggests that tau can adopt disease-specific pathological conformations that spread through the nervous system by a prionic mechanism to cause diseases like Alzheimer’s disease (AD) and other tauopathies (35–38). According to this model, structurally distinct tau conformers replicate by inducing the templated conversion of native tau proteins into pathological forms (“seeding”) and produce predictable patterns of pathology, akin to prion strains. Faithful propagation of abnormal tau has been demonstrated in both cell and mouse models (39–45). Insight into the precise nature of different tau strains and their association with different tauopathies could lead to a better understanding of the molecular mechanisms underlying disease as well as the development of improved diagnostics and therapies (46). Recent work has shown that Guam ALS/PDC brain extracts display tau seeding activity in fluorescence resonance energy transfer (FRET)-based biosensor cells (47). Additionally, electron cryo-microscopy (cryo-EM) analysis of tau filaments isolated from ALS/PDC nervous tissues has shown that abnormal tau in these cases is structurally similar or identical to the tau found in chronic traumatic encephalopathy (CTE), a disease associated with repetitive brain trauma (48, 49). This particular tau fold has thus far also been observed in subacute sclerosing panencephalitis (SSPE), a rare consequence of measles virus infection (50), and in vacuolar tauopathy (VT), a form of dementia caused by mutation in the gene encoding valosin-containing protein (*VCP*) (51). The tau amyloid structure itself appears to encode information required to establish unique patterns of disease and it will therefore be of considerable interest to further explore the characteristics of conformationally diverse disease-associated tau species.

In the present study, we used an alanine mutational scanning (Ala scan) approach (52) to determine the seeding signature of tau in Guam ALS/PDC cases. This strategy relies on a prion species barrier concept for tau, i.e., the notion that specific abnormal tau assemblies cannot serve as a template for native tau monomers if the amino acid sequences are incompatible. By sequentially mutating each residue in the repeat domain (RD) of tau (corresponding to the core of the tau protein that is responsible for its aggregation) to a functionally neutral alanine, an incorporation profile of tau can be established through cytometry-based seeding experiments. It has been demonstrated that Ala scan can robustly discriminate between different tau strains (52, 53). We find that, when compared to other tauopathy profiles, ALS/PDC tau seeding profiles resemble those of CTE-type tau. This is in agreement with the reported near-atomic level cryo-EM structures of ALS/PDC tau (48). Thus, the data supports the structure-based classification of Guam ALS/PDC as a CTE-like tauopathy.

## Results

### FRET biosensor cells detect tau seeds in ALS/PDC brain and spinal cord tissues

We included 14 ALS/PDC cases (Age: 61.07, SD 9.75; M:F 2.5:1) as well as 8 control cases (Age: 52.75, SD 16.46; M:F 3:1), the latter corresponding to Guam subjects that did not develop symptomatic disease. An overview of all tissue samples used in this study can be found in **Table 1** and the **Supplementary Information** (this overview also includes identifiers for associated whole-genome sequencing and RNA-sequencing data on these nervous tissues).

**Table 1:** Sample IDs used in the study and average seeding percentage in 3R3R, 3R4R and 4R4R biosensors.

|  |  |  |  |  |  |  |  |
| --- | --- | --- | --- | --- | --- | --- | --- |
| Case # | Pathology Diagnosis | Age | Gender | Tissue Type | 3R3R Seeding (%) | 3R4R Seeding (%) | 4R4R Seeding (%) |
| 1 | ALS/PDC | 69 | F | Frontal Cortex | 0.01 | 0.89 | 6.86 |
| 2 | ALS/PDC | 61 | M | Frontal Cortex | 0.05 | 0.06 | 0.14 |
| 3 | ALS/PDC | 52 | F | Frontal Cortex | 0.56 | 1.89 | 5.64 |
| 7 | ALS/PDC | 71 | M | Spinal Cord | 0.02 | 1.30 | 2.35 |
| 9 | ALS/PDC | 60 | M | Frontal Cortex | 0.06 | 0.14 | 0.26 |
| 10<sup>(1)</sup> | Control | 72 | M | Frontal Cortex | 0 | 0 | 2.42 |
| 11<sup>(1)</sup> | Control | 72 | M | Spinal Cord | 0 | 0 | 2.21 |
| 12<sup>(2)</sup> | ALS/PDC | 54 | M | Frontal Cortex | 0.03 | 1.41 | 4.59 |
| 13<sup>(2)</sup> | ALS/PDC | 54 | M | Spinal Cord | 0 | 0 | 3.86 |
| 14<sup>(3)</sup> | ALS/PDC | 64 | F | Frontal Cortex | 0 | 0 | 1.33 |
| 15<sup>(3)</sup> | ALS/PDC | 64 | F | Spinal Cord | 0.08 | 0 | 17.15 |
| 17 | ALS/PDC | 73 | M | Spinal Cord | 0 | 0 | 8.63 |
| 19 | ALS/PDC | 51 | F | Frontal Cortex | 0.19 | 0.99 | 3.69 |
| 24 | Control | 68 | F | Spinal Cord | 0 | 0 | 0 |
| 25<sup>(4)</sup> | Control | 57 | F | Frontal Cortex | 0 | 0 | 2.22 |
| 26<sup>(4)</sup> | Control | 57 | F | Spinal Cord | 0.06 | 0 | 0.75 |
| 28 | Control | 48 | M | Spinal Cord | 0 | 0 | 0 |
| 29<sup>(5)</sup> | ALS/PDC | 49 | M | Frontal Cortex | 0 | 0 | 0 |
| 30<sup>(5)</sup> | ALS/PDC | 49 | M | Spinal Cord | 0.07 | 0 | 0 |
| 31 | Control | 31 | M | Frontal Cortex | 0 | 0 | 0 |
| 32 | Control | 27 | M | Frontal Cortex | 0.16 | 0 | 0 |
| 35 | Control | 63 | M | Spinal Cord | 0 | 0.01 | 0.14 |
| 36<sup>(6)</sup> | ALS/PDC | 61 | M | Frontal Cortex | 0 | 0.26 | 6.60 |
| 37<sup>(6)</sup> | ALS/PDC | 61 | M | Spinal Cord | 0 | 0.04 | 17.0 |
| 38<sup>(7)</sup> | ALS/PDC | 76 | M | Frontal Cortex | 0 | 0.01 | 5.62 |
| 39<sup>(7)</sup> | ALS/PDC | 76 | M | Spinal Cord | 0 | 0.01 | 3.33 |
| 40<sup>(8)</sup> | ALS/PDC | 69 | M | Frontal Cortex | 0 | 0.56 | 2.97 |
| 41<sup>(8)</sup> | ALS/PDC | 69 | M | Spinal Cord | 0 | 0.03 | 7.16 |
| 42<sup>(9)</sup> | ALS/PDC | 45 | M | Frontal Cortex | 0 | 0 | 1.68 |
| 43<sup>(9)</sup> | ALS/PDC | 45 | M | Spinal Cord | 0.03 | 0 | 0 |
| 44<sup>(10)</sup> | Control | 56 | M | Frontal Cortex | 0 | 0 | 0.65 |
| 45<sup>(10)</sup> | Control | 56 | M | Spinal Cord | 0 | 0 | 0 |

<sup>(N)</sup> Sample IDs labelled with a <sup>(N)</sup> represent provenance from the same case. Abbreviations: F, Female; M, Male

To measure tau seeding activity in Guam ALS/PDC and control nervous tissues, we used engineered tau biosensor cell lines that carry constructs genetically encoding the 3-repeat (3R) or the 4-repeat (4R) wild-type tau RD (residues 246-408) fused to the fluorophores Cerulean and Ruby. Three biosensor cell lines were used, one consisting of 3R tau only (3R/3R), one consisting of 4R tau only (4R/4R), and one consisting of a mixture of 3R and 4R tau (3R/4R) (**Fig. 1**). If tissue extract containing seed-competent tau is added to these cells, this will lead to a FRET signal due to incorporation of tau into seeded aggregates, which can be detected by flow cytometry. Samples can exhibit preferential seeding on either the 3R/3R, 3R/4R or 4R/4R biosensors, presumably depending on the isoform composition of the tauopathy homogenates.

**Figure 1:**
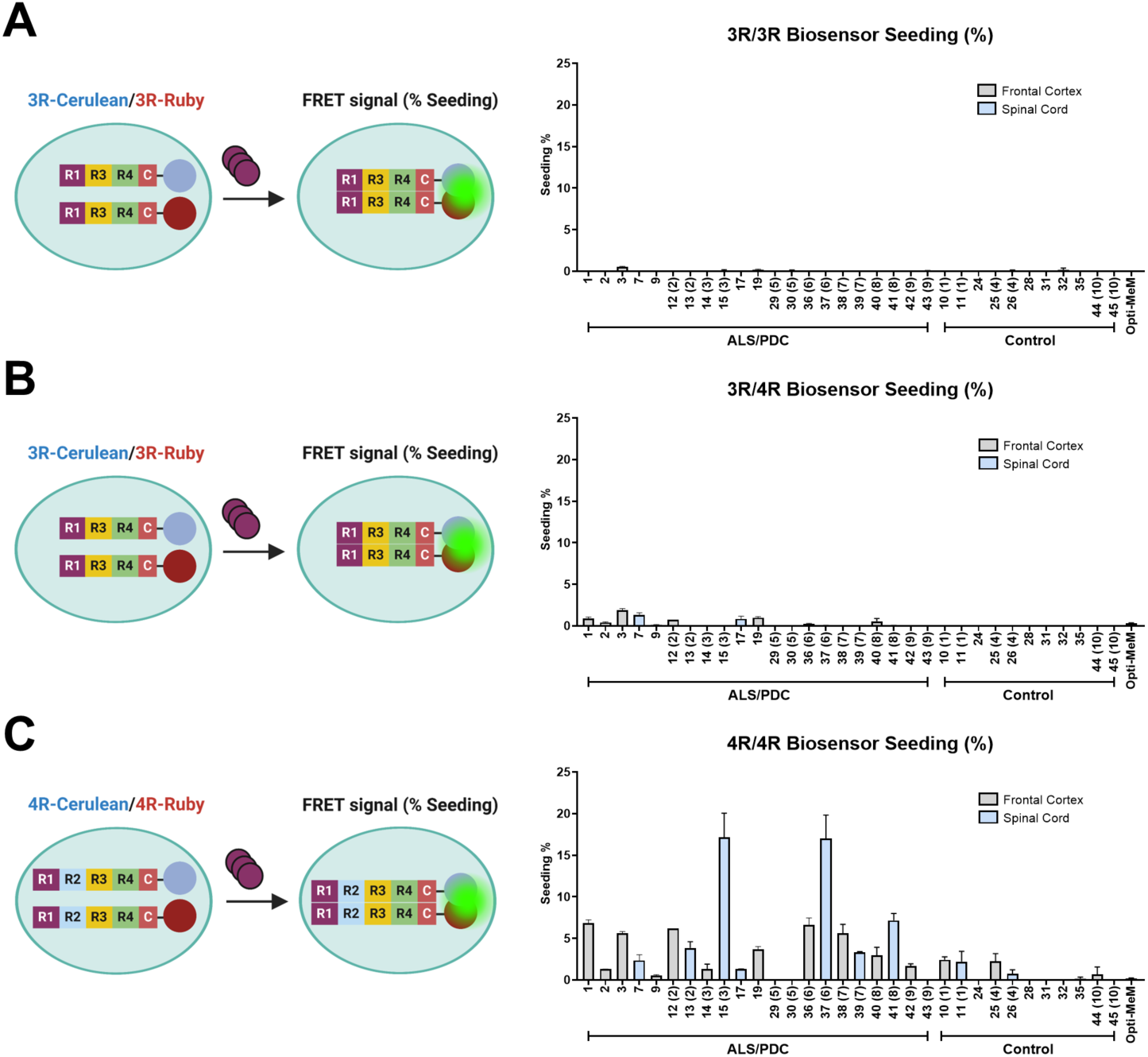
Tau seeds in Guam ALS/PDC brain and spinal cord tissues. Schematic representations and outcomes of the tau seeding assays. ALS/PDC nervous tissue extracts were seeded onto biosensor cells containing either 3-repeat (3R) or 4-repeat (4R) tau fused to different fluorophores (Cerulean, Ruby). The three cell lines used harbored 3R tau only (3R/3R), 3R and 4R tau (3R/4R), and 4R tau only (4R/4R). Cells were fixed 72 h post-seeding and evaluated for FRET by flow cytometry. 3.5 μg of nervous tissue homogenate material from ALS/PDC cases and Guam controls (Control) were used for the seeding assay. The three cell lines (3R/3R [**A**], 3R/4R [**B**] and 4R/4R [**C**]) were seeded with the homogenates. Quantification of seeding efficiency was made as the % of cells exhibiting FRET signal (Seeding %).

We incubated the three biosensor cell lines with brain homogenate from the ALS/PDC cases and controls. We observed that tau seeding of ≥2% could be detected in the 4R/4R biosensors (**Fig. S2**), but not the 3R/3R or 3R/4R biosensors, in the brain (frontal cortex) and spinal cord of a subset of ALS/PDC cases (n=10/14). Tau seeding was detected in 7/12 frontal cortex samples, and 7/9 of the spinal cord samples. Interestingly, select Guam control cases (n=2/8) also showed seeding activity in the biosensor cells, with a similar preference for the 4R/4R biosensors (**Table 1**, **Fig. 1**). Of note, the tau seeding activity may have been limited by tissue availability, as only 3.5 μg of tissue per well in a 96-well plate were used for seeding in the biosensors.

The presence of tau seeds in ALS/PDC brain (frontal cortex) is in agreement with published data (47). To our knowledge, this is the first demonstration of seeding-competent tau in the spinal cord. It should be noted that the mean age of the included Guam control cases (mean age: 52.75) was lower than that of ALS/PDC cases (mean age: 61.07), which may have impacted the outcome of the seeding assays. It is unknown what causes the preference for seeding in the 4R/4R tau biosensors in these cases (ALS/PDC is recognized as a 3R + 4R tauopathy), but the presence of specific 4R tau species that are more seed-competent or a switch in isoform usage could underlie this effect (52, 54, 55).

Overall, the highest tau seeding activity among all cases tested was found in spinal cord tissue, samples 15 and 37. A case used for previous cryo-EM analysis of tau (sample 17), in which ALS/PDC tau filaments were found to adopt the CTE fold (48), also showed seeding activity. The cases that had ≥2% seeding activity in the seeding reporter cells were selected for tau Ala scan experiments.

### Alanine mutational scanning of ALS/PDC tau identifies a CTE-like signature

We next exploited a tau Ala scan functional genetics approach (52) to determine the nature of tau seeds in Guam ALS/PDC. By sequentially introducing alanine substitutions into the tau RD (residues 246-408), the ability of the tau point mutants to incorporate into unique strains will be differentially affected (**Fig. 2A**). Identifying the residues required to form each strain’s core allows one to “fingerprint” tau strains and precisely distinguish conformationally different species (52, 53).

**Figure 2:**
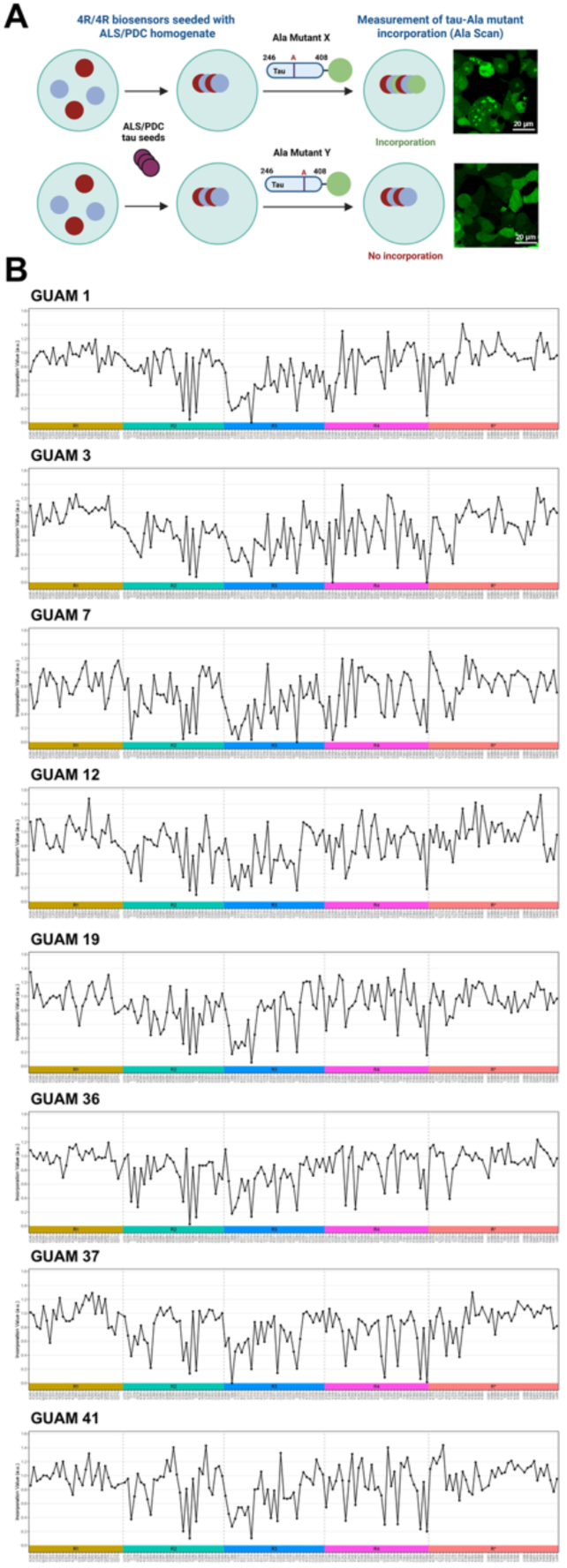
Tau Ala scan of ALS/PDC samples. **A**. Tau Ala scan overview. The 4R/4R tau biosensor cells were seeded with homogenate from the ALS/PDC cases. After seeding occurred, a lentiviral library of tau Ala point mutants spanning the tau repeat domain (RD) conjugated to fluorophore Clover (i.e. X, Y) was added to the seeded biosensor cells, along with the appropriate controls (wild-type tau RD and I277P-I308P double proline tau RD). Incorporation of each point mutant into the seeded aggregates is measured as the FRET signal between Clover in the Ala point mutants and Cerulean in the cell aggregates. The incorporation profile fingerprint can faithfully discriminate between different tau strains. Figure adapted from (52). **B**. ALS/PDC brain homogenates from samples #1, #3, #7, #12 (left), #19, #36, #37, and #41 (right) were incubated with the tau 4R/4R biosensor cells. After a 48-hour incubation period that allowed inclusion formation, the biosensor cells were re-plated in triplicates and incubated with the tau-(246-408)-Ala-mEOS3.2 mutant lentivirus library. The cells were harvested 72 hours after transduction. The resulting tau Ala scan is shown for each sample. The incorporation value (a.u.) represents the FRET median fluorescence intensity between Clover and Cerulean for each position normalized by the positive (WT-tau RD-mEOS3.2) and negative controls (I277P-I308P-tau RD-mEOS3.2). Each position in the tau RD is represented on the X axis. Positions K274A, T386A and P397A were excluded from the analysis due to failure of transduction.

We incubated Guam ALS/PDC nervous tissue extracts with the mutant tau biosensor cells and through flow cytometry analysis were able to extract an incorporation profile (**Fig. 2B**). Samples 36 and 37 originated from the same patient, and corresponded to the frontal cortex and spinal cord, respectively. When we compared the Ala scan profiles from these samples with those of other tauopathies, we found that all Guam samples evaluated (n=8, **Table 2**) most closely resembled the profiles seen in CTE with an average R^2^ (R^2^ Avg) of 0.566 and a SD of 0.037 (**Fig. 3**). Observed correlations against CTE were significantly better than those against AD (R^2^ Avg 0.294 +/- SD 0.044), progressive supranuclear palsy (PSP) (R^2^ Avg 0.165 +/- SD 0.039), and corticobasal degeneration (CBD) (R^2^ Avg 0.226 +/- SD 0.054). We could not extract a profile for all cases that exhibited seeding due to limited tissue availability. When the 8 ALS/PDC profiles were averaged, the mean ALS/PDC profile strongly correlated with CTE (R^2^ 0.805) (**Fig. S3**).

**Figure 3:**
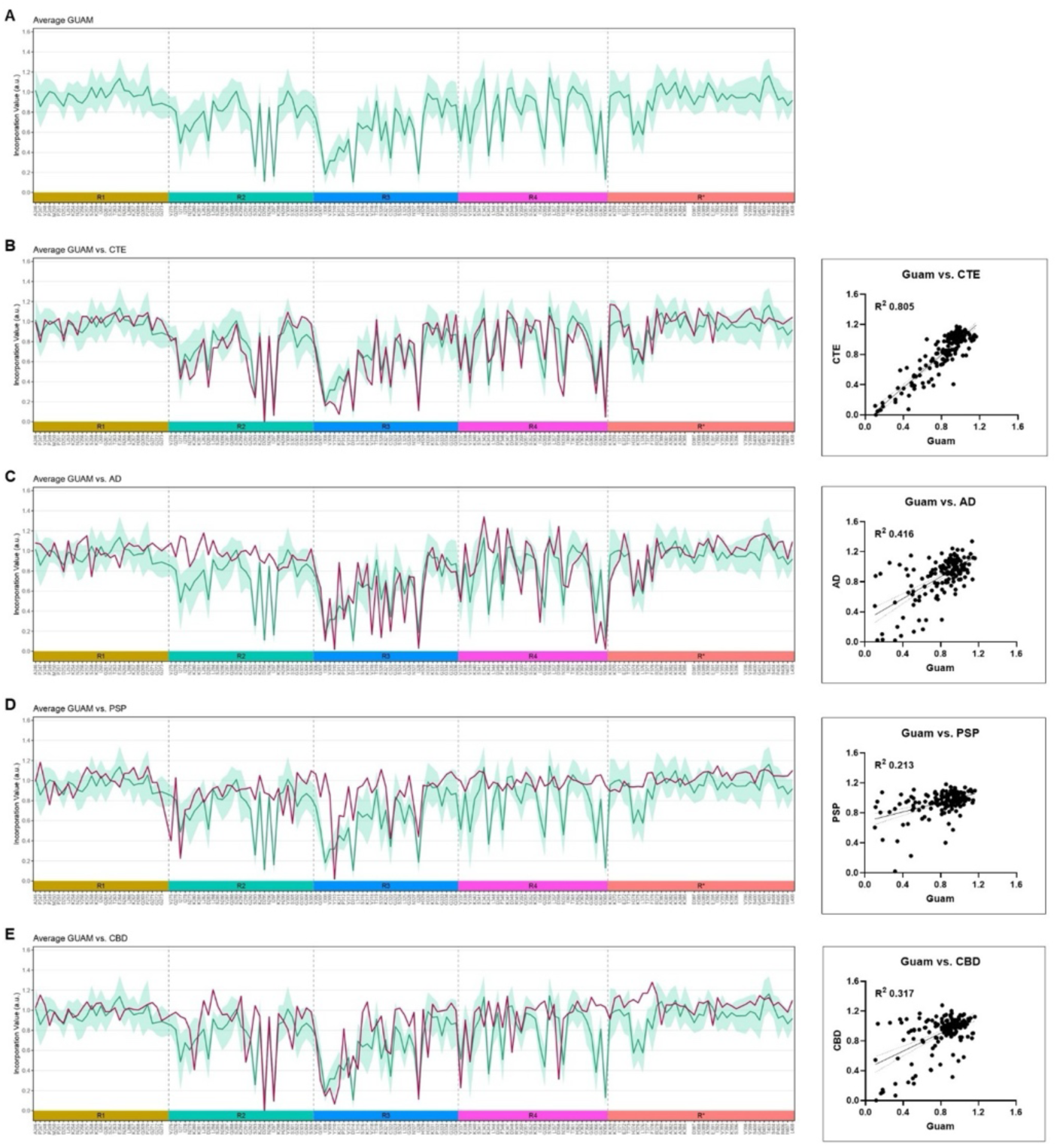
Average incorporation profile for ALS/PDC and comparison with other tauopathy average profiles. **A**. Mean incorporation profile (line) for the ALS/PDC samples (n=8). The standard deviation (SD) for each position is represented with the teal-colored area. **B**, **C**, **D**, **E**. Comparison of the average ALS/PDC incorporation profile with the average CTE (n=2), AD (n=8), PSP (n=5) and CBD (n=8) profiles. The best correlation was observed with CTE (R^2^ 0.805).

**Table 2:** Sample IDs used in the 4R/4R Alanine Scan.

| Table 2: Sample IDs used in the 4R/4R Alanine Scan |  |  |  |  |
| --- | --- | --- | --- | --- |
| Case # | Pathology Diagnosis | Age | Gender | Tissue Type |
| 1 | ALS/PDC | 69 | F | Frontal Cortex |
| 3 | ALS/PDC | 52 | F | Frontal Cortex |
| 7 | ALS/PDC | 71 | M | Spinal Cord |
| 12 | ALS/PDC | 54 | M | Frontal Cortex |
| 19 | ALS/PDC | 51 | F | Frontal Cortex |
| 36<sup>(6)</sup> | ALS/PDC | 61 | M | Frontal Cortex |
| 37<sup>(6)</sup> | ALS/PDC | 61 | M | Spinal Cord |
| 41 | ALS/PDC | 69 | M | Spinal Cord |

<sup>(N)</sup> Sample IDs labelled with a <sup>(N)</sup> represent provenance from the same case.

Cluster analysis confirmed that the Guam ALS/PDC samples show a comparable incorporation profile to CTE i.e., the Guamanian samples clustered together with CTE, when compared to profiles from other tauopathies (**Fig. 4A**). Importantly, the data indicated that ALS/PDC tau seeds are different from AD-type tau seeds, which are known to be structurally similar to CTE-type tau, but have a slightly different (i.e., less “open”) conformation (48, 49). In this analysis, we also included two primary age-related tauopathy (PART) samples, which we had previously found to match the incorporation profile of AD (53). Profiles of some ALS/PDC samples exhibited poor correlation amongst each other (i.e., sample 7 vs. sample 19; sample 3 vs. sample 41), despite exhibiting a robust correlation with the CTE profile (**Fig. 4B**). However, we deem this effect may have been confounded by low seeding activity due to limited tissue availability in these samples, which contributed to an increased background signal in non-residue hit positions.

**Figure 4:**
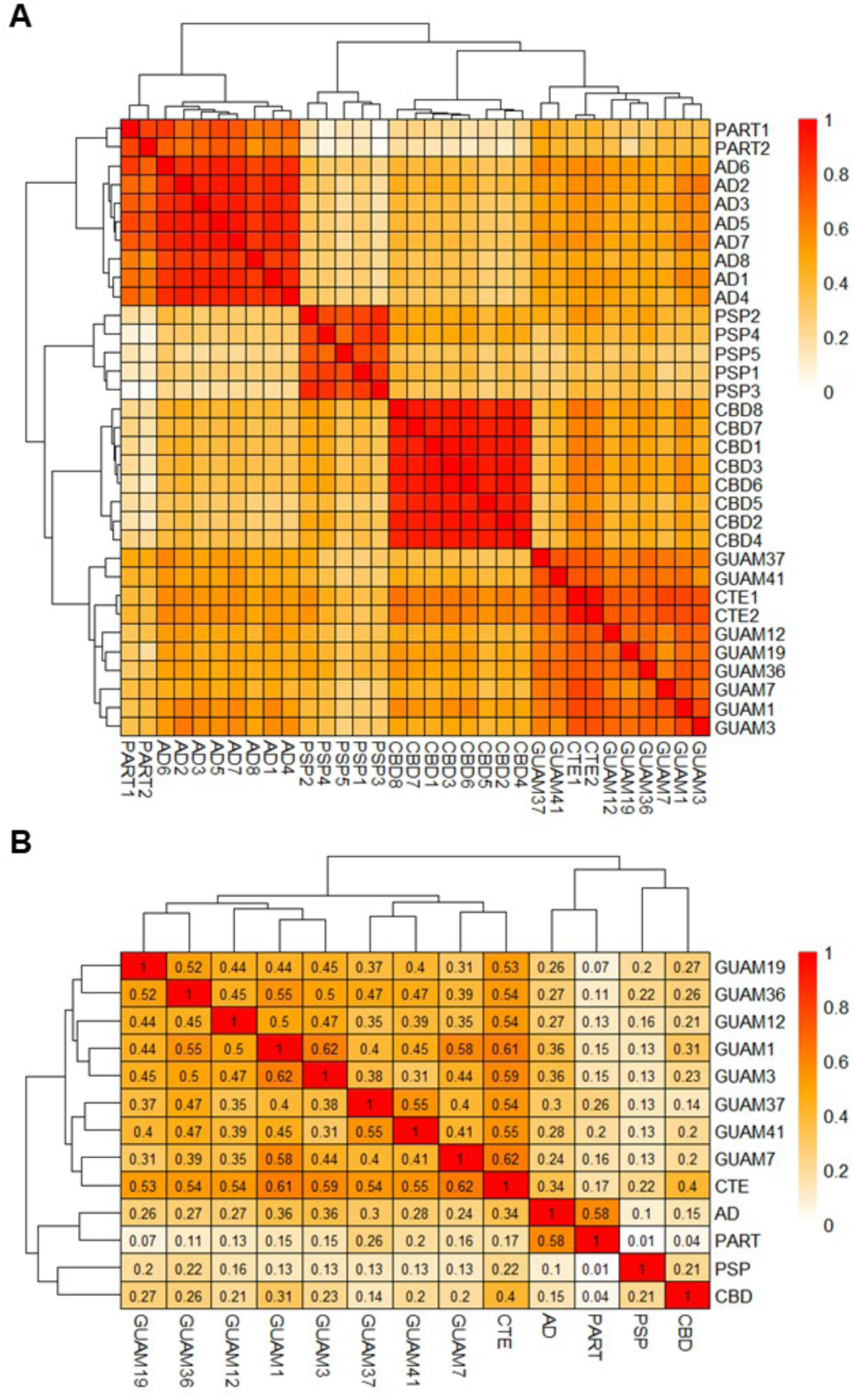
Clustered heatmap analysis of ALS/PDC and other tauopathies. **A**. Clustered heatmap of ALS/PDC, AD, CTE, CBD, PSP and PART samples. The ALS/PDC samples (GUAM) clustered together with the CTE cases. **B**. Clustered heatmap analysis of ALS/PDC samples and the average profiles for AD (n=8), CTE (n=2), CBD (n=8), PSP (n=5), and PART (n=2). The average correlation between all ALS/PDC samples was R^2^ 0.44 +/- SD 0.078.

Our data demonstrates that ALS/PDC tau seeds closely resemble those present in CTE. We therefore propose to classify ALS/PDC as a CTE-type tauopathy.

## Discussion

We report here on the seeding behavior of tau in Guam ALS/PDC, a neurodegenerative disorder of unknown cause that was previously highly prevalent in the Pacific Island of Guam (Mariana Islands). In agreement with a recent publication (47), we found that Guam ALS/PDC brain samples display robust tau seeding activity in tau biosensor cells. Notably, we also detected strong tau seeding activity in spinal cord tissue. To our knowledge, this is the first description of seed-competent tau in spinal cord. The significance of the presence of tau seeds in the spinal cord region is not clear currently, however, tau inclusions in the spinal cord appear to be present in a number of neurological disorders other than Pacific ALS/PDC (23, 56–60). Spinal cord tau pathology may be more widespread and perhaps defines a subset of disease cases. Tau pathology in the spinal cord was found to induce tactile deficits and cognitive impairment in AD via dysregulation of cholecystokinin-expressing neurons (61). A comprehensive analysis of tau pathology in spinal cord tissues across neurodegenerative diseases, including measurements on tau seeding activity, would provide additional context. Further, we found that 2 out of 8 Guam non-neurological control subjects also showed seeding activity in the biosensor cells. This is in keeping with seeding data from another study (47) and with previous neuropathological findings in Guamanians, i.e., tau pathology appears to be a frequent background feature in the Guam population (32, 33). A plausible explanation is that these tissues represent presymptomatic cases. An alternative, and perhaps more provocative, explanation is that some individuals may possess resilience against tau seeds (47). Another possibility is that the tau seeds detected in our assay may not be a dominant factor in determining disease phenotypes overall. Tau seeds in all evaluated tissues showed a strong preference for the 4R/4R tau biosensors. This preference was also observed in the aforementioned study on tau seeds in Guam ALS/PDC (47). CTE tau appears to also seed preferably onto 4R/4R biosensors (52). It is not clear why Guam ALS/PDC and CTE, which are both 3R + 4R tauopathies, feature tau seeds that prefer to seed on 4R-only tau reporter cells, but a potential explanation is that tau seeds behave like this due to their specific structural conformation. A switch in isoform usage could also account for the observed preference for seeding onto the 4R biosensors (52, 54, 55). Distinct tau conformers may be present within individuals and specific 4R tau species in ALS/PDC could have particularly strong seeding activity. The cellular source from which these tau seeds arise in ALS/PDC is not known.

We selected 8 samples that displayed robust seeding activity in the 4R/4R biosensors (≥2% seeding threshold) and had sufficient tissue availability to perform tau Ala scan assays. We found that all samples analyzed showed an Ala scan incorporation profile that most closely resembled that of CTE-type tau. A consensus ALS/PDC incorporation profile robustly correlated with CTE (R^2^ 0.805). In accordance with our findings, *Qi et al.* recently reported on the cryo-EM structures of tau filaments in ALS/PDC brain and spinal cord tissues (48), showing that tau filaments in ALS/PDC adopt folds that appear to be identical to those identified in CTE (49), SSPE (50) and VT (51), disorders that are associated with repetitive brain trauma, measles virus infection, and *VCP* mutation, respectively. Cryo-EM has allowed the development of a structure-based classification of tauopathies (36, 62) and provides a basis for categorizing ALS/PDC tau filaments. Like in CTE, two types of tau filaments (Type I and Type II) were present in variable amounts, each with an unknown, internal density in the β-helix of the structured core (48). The unknown density may represent a cofactor for filament formation. Determining the nature of this extra density is not straightforward, because amyloids extracted from human brains contain different types of contaminants that make finding out what specific densities are present difficult. The ALS/PDC (/CTE) tau fold differs from the AD tau fold, because of the more open conformation of the β-helix region. Type I and Type II filaments are both made of two identical protofilaments, with different interprotofilament packing, i.e., Type I and Type II filaments are ultrastructural polymorphs. A third type of tau filament (Type III), in which two protofilaments pack with opposite polarities, was also identified in both CTE and ALS/PDC (48). The classification of ALS/PDC as a CTE-type tauopathy is now further supported by the Ala scan data in this study, which identified a hit pattern closely matching CTE but not AD, PSP or CBD. Interestingly, tau filaments extracted from Kii ALS/PDC tissues had identical structures on cryo-EM (48), providing support for the notion that ALS/PDC of Guam and Kii are similar or identical diseases. No Kii ALS/PDC cases were included in the present study, but it would be of interest to analyze tau from those cases in the Ala scan assay in the future. Understanding the cause of ALS/PDC in Guam may provide insight into the cause of Kii ALS/PDC and vice versa. No neuropathological studies are available for the disease cluster in Papua, rendering comparisons of pathology features with that geographic focus of disease impossible at this time. Ala scan of other tauopathies that adopt the CTE fold, i.e., SSPE and VT, may also reveal similar incorporation profiles. We anticipate that Ala scan of diverse tauopathies will result in a classification scheme that is compatible with the classifications established by cryo-EM, because the tau amyloid structures are coupled to its seeding behavior. This is particularly important for rare tauopathies, such as ALS/PDC, in which tissue availability constitutes a limiting factor and cryo-EM may not be feasible. In this study, we were able to extract a profile for all seeding cases where 1 mg of material was available.

Shared structural features of tau across different disease conditions suggest a possible commonality in disease mechanisms. Previous studies identified alterations related to several processes, such as neuroinflammation (together with vascular changes and indications of blood-brain barrier impairment) (63, 64), mitochondrial dysfunction and oxidative stress (65–67), RNA metabolism (68, 69), and protein homeostasis (proteostasis) pathways (70) (**Fig. S4**) in ALS/PDC tissues. Of these processes, neuroinflammation is of particular interest with regard to CTE-type tauopathy, as a pathogenic role for neuroinflammatory changes has been recognized in brain injury and CTE (71–74). Neuroinflammation has been increasingly associated with neurodegeneration and tauopathy (75–77). It is possible that chronic neuroinflammation plays a role in the assembly of abnormal tau (e.g., local inflammation in specific neuroanatomic regions could give rise to specific tau folds) or that tau pathology contributes to inflammatory changes. Neuroinflammation may also modify tauopathy, e.g., by altering the spread of tau or by affecting neuronal health and tissue functional decline. Tauopathies characterized by the CTE fold may converge on similar disease mechanisms, regardless of the initial insult or trigger. We speculate that there may be mechanistic overlap between ALS/PDC and other CTE-like tauopathies and nominate neuroinflammation as a candidate mechanism. The precise roles of neuroinflammation and other mechanisms in ALS/PDC pathogenesis remain to be explored further (under investigation).

Various aspects of ALS/PDC tau seeds remain unclear. For example, although there currently is no compelling evidence for ALS/PDC being transmissible [i.e., attempts to show that ALS/PDC can be transmitted have been unsuccessful (78, 79)], it is not known whether ALS/PDC tau seeding is attainable in animal models. It will be important to establish whether ALS/PDC tau is seed-competent in intact animals. Further, tau seeding in CTE has been examined in relation to tau deposition during disease progression (80), but it is not clear whether tau seeding parallels disease severity in ALS/PDC. A detailed and reliable neuropathological staging of tau lesions in Guam ALS/PDC cases is not available, which has precluded such an analysis. The determinants of specific tau folds in patient tissues are incompletely understood and likely involve different factors, such as tau sequence and isoform composition, post-translational modifications and tau-interacting molecules. It seems probable that the pattern and distribution of tau pathology are connected to specific cellular environments that influence tau aggregation behavior. It has been suggested that, in addition to the structure of the ordered tau core, phosphorylation in the “fuzzy coat” (i.e., the disordered regions of tau on both the C-terminal and N-terminal sides of the ordered filament core) confers the seeding capacity of different tau filaments (81, 82). It will be of interest to map in detail the post-translational modifications, including phosphorylation sites, of ALS/PDC tau proteoforms. Previous work has identified several phosphorylated epitopes on Guam ALS/PDC tau (47, 83). A preliminary analysis of Guam ALS/PDC tau phosphorylation identified similar modifications (**Fig. S5**), but more rigorous methods should be applied to explore such modifications comprehensively (84–86). The parameters that govern tau prion-like replication, spread, and clearance, as well as tau aggregation behavior in general (87), merit further study. Special attention should be given to factors known to play a role in tau proteostasis regulation, such as VCP(/p97) (88, 89), because they may affect aggregate processing and underlie selective vulnerability in disease. To date, mutation in *VCP* is the only genetic link to the CTE tau fold (51).

It is important to emphasize that ALS/PDC is a multiproteinopathy that does not only involve tau deposits, but also features other protein abnormalities, such as pTDP-43 inclusions (17–19, 90) (**Fig. S1**). It has been put forward that Guam ALS/PDC is, in fact, a multi-prion disorder, that involves tau prions as well as other proteopathic seeds, i.e., Aβ prions (47). The detection of Aβ seeds in ALS/PDC brains is somewhat remarkable given the relative absence of extracellular Aβ plaques (91–93) [although a number of studies do report on Aβ deposits in ALS/PDC (90, 94–97), and senile plaques may be more frequent in subjects of advanced age]. Cryo-EM analyses of Guam ALS/PDC cases also found Type II Aβ42 filaments in one tissue sample (48). One possibility is that certain protein seeds can be present in the absence of evident deposits of such proteins on histology (47, 98–101). Attempts to detect α-synuclein seeds in ALS/PDC using biosensor assays were unsuccessful (47), but the most likely brain regions to contain such seeds, i.e., amygdala and cerebellum, were not included in those experiments. Another study did find evidence for α-synuclein seeds in Guam ALS/PDC spinal fluid through seed-amplification assays (102). Methodological differences could underlie these discrepancies. Sampling from different nervous system regions may be required to adequately detect certain seed-competent proteins. Also, select seeds may only be present in a subset of cases. We focused in our experiments on tau seeding, but future studies should investigate the role of other protein seeds [e.g., TDP-43 seeds (103)] in more detail, also taking into consideration potential interactions between various pathologies. This will also be important to establish what factors differentiate ALS/PDC from other pathologies.

Several limitations should be pointed out. First, the number of samples and amounts of tissue included for this study were low. Because of limited availability of tissues, we relied on archival frozen tissue specimens for these experiments. This also meant we were restricted to a limited number of neuroanatomical regions (i.e., frontal cortex and spinal cord). These limitations may contribute to variation observed in our assays. In addition to the low amount of input material available for our assays, differences in sampling and sample processing may have contributed to overall noise in the data. The variation between samples could also be related to the heterogeneity of ALS/PDC itself, which shows considerably variety in presentation. Possibly, different tau conformers, with similar or distinct immunophenotypes, are present within individual cases. Postmortem studies are confined to describing processes at work at the end stage of disease and cannot be used to establish the relative timing of molecular alterations that occur during the disease course. The cells that persist at the end stages of disease may be resilient to disease-associated processes, which could lead to bias. While seeded aggregation in the biosensor cells varies with the position of alanine mutations in the tau sequence, in a tau fold-dependent manner, care should be taken in interpreting the Ala scan results in terms of input seed structures. The structure of the seeded aggregates formed in the biosensors could not be determined due to the fluorescent protein tags hampering cryo-EM analysis.

In summary, in this study we leveraged Ala mutational scanning to determine the incorporation profile of Guam ALS/PDC tau into aggregates. We found that Guam ALS/PDC tau showed an Ala scan incorporation profile that is similar to that of CTE-type tau. Based on this data, and on the cryo-EM structures of ALS/PDC tau as well as observations regarding the regional distribution of tau pathology in ALS/PDC (e.g., laminar selectivity), we suggest to classify ALS/PDC as a CTE-like tauopathy. The findings support the use of the Ala scan assay for rapid and unbiased classification of tauopathies, especially when tissue availability is limited and cryo-EM cannot be performed. Further optimization of the assay (104), as well as inclusion of other neuroanatomic regions, will help better characterize these disorders.

## Materials and Methods

### Human Brain Samples

Post-mortem brain and spinal cord tissue specimens from Guam ALS/PDC cases and Guam controls were used for experiments in this study. Use of human postmortem tissues was approved by the Ethics Committee of Niigata University [2020–0019].

### Tissue Processing

Frozen brain tissue samples were suspended in Tris-buffered saline (TBS) containing cOmplete mini protease inhibitor tablet (Roche) at a concentration of 10% weight/volume in a 1.5 ml low-binding tube (Thermo Fisher Scientific). The tissue was homogenized using a handheld homogenizer (BT Lab Systems) for 1 minute in a vented hood. The homogenates were centrifuged at 21,000xg for 20 minutes. The supernatant was collected as the total soluble protein lysate. Protein concentration was measured using a Pierce 660 Assay (Pierce) and the concentration was adjusted to 1 mg/ml. The fractions were aliquoted into low-binding tubes (Thermo Fisher) and frozen at -80 ^°^C until further use. Prior to use in the seeding assays, the thawed homogenates were sonicated for 1 minute using a Q500 Sonicator (QSonica).

### Tau RD Biosensor Cell Lines

For this study, the following wild-type (WT) human tau repeat domain (RD) HEK293T biosensor cell lines were used for tau seeding assays:

#### Tau RD 3R/3R Cells

3R/3R cells harbor the 3-repeat (3R) version of WT human tau RD (residues 246-408) C-terminally fused to mCerulean (Cer) and mRuby (Ruby) fluorescent protein.

#### Tau RD 3R/4R Cells

3R/4R cells harbor the 3R version of WT human tau RD (246-408) C-terminally fused to Cer, and 4R tau fused to Ruby.

#### Tau RD 4R/4R Cells

4R/4R cells harbor 4R tau fused to Cer and Ruby.

### Tau Seeding Assays

Tau RD 3R/3R, 3R/4R and 4R/4R cells were plated into 96-well plates at a density of 20,000 cells per well. After 18 hours, the cells were transduced with a complex of 0.75 µl of lipofectamine 2000 (Invitrogen), 9.25 µl Opti-MEM (Gibco), and 3.5 µg of 10% weight/volume brain homogenate, for a final treatment volume of 13.5 µl per well. The brain homogenate volume was limited by ALS/PDC tissue availability. After 72 hours, the cells were harvested using 0.25% trypsin-EDTA (Gibco) digestion for 5 minutes at 37°C, followed by quenching with Dulbecco’s Modified Eagle Medium (DMEM, Gibco), and transferred to 96-well U bottom plates. The samples were then centrifuge for 5 minutes at 1500 rpm, and the pellet was resuspended and fixed in 2% paraformaldehyde (PFA) in Phosphate-buffered saline (PBS, Gibco) for 10 minutes. The cells were centrifuged again, and the pellet resuspended in 150 μl of PBS for flow cytometry analysis.

### 4R/4R Alanine Scan

For the alanine scan incorporation assay, 4R/4R biosensor cells were plated at a density of 20,000 cells per well into two 96-well plates per case. This was then followed by treatment with the sonicated homogenates for selected ALS/PDC cases, as described above. 48 hours after initial seeding, once seeded aggregates were observed in the biosensor cells by microscopy imaging, the cells on each plate were re-plated into three 96-well plates to generate technical replicates. The cells were then treated with a library of lentivirus harboring tau RD alanine point mutants spanning from positions 246 to 408, and C-terminally conjugated to mEOS3.2. One alanine point mutant construct was added per well, in addition to the appropriate positive (WT tau RD), and negative controls (double proline I277P-I308P tau RD). 72 hours after lentiviral transduction, the cells were harvested and fixed in PFA as described above.

### Flow Cytometry Analysis

The Attune CytPix Flow Cytometer (Thermo Fisher) was used to perform Fluorescence Resonance Energy Transfer (FRET) analysis. The biosensor cells were analyzed by gating for homogeneous side-scatter and forward-scatter. The FRET signal between Cer and Ruby was measured in the Pacific Blue and Qdot605 channels with gating for cells that contained seeded aggregates in the different biosensors (3R/3R, 3R/4R, 4R/4R). For the alanine scan analysis, a subsequent gate was used for cells positive for FITC (non-photoconverted mEOS3.2), which corresponded to seeded cells that had been successfully transduced with the tau RD-Ala-mEOS3.2 point mutants. Within this population, the FRET signal between Cer and mEOS3.2 was measured in the Pacific Blue and AmCyan channels, and a narrow gate of bright cells was drawn to calculate the median fluorescence intensity (MFI) in the AmCyan channel. The MFI of this FRET signal was used as an indicator of the degree of incorporation of each tau RD alanine point mutant into the seeded aggregates in the 4R/4R biosensors. Alanine mutants K274A, T386A, and P397A were excluded from the scan due to low transduction efficiency.

### Alanine Scan Incorporation Data Analysis

The FRET MFI values between Cer and mEOS3.2 measured in the AmCyan channel were normalized by plate to prevent batch variation. The tau RD alanine mutants in the N-terminus and the C-terminus of the sequence do not affect incorporation in our assays. Thus, we used the first twenty mutants in the N-terminus, and the last ten mutants in the C-terminus, as well as the minimum MFI in each plate, to normalize the data and obtain the Incorporation Ratio for each residue position. We used the following formula to obtain this:

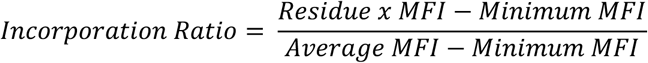

In this formula Residue x MFI is the FRET MFI in AmCyan for each position, Minimum MFI is the minimum FRET value in the scan for each plate, and Average MFI corresponds to the Average MFI for the first 20 residues (on the first plate), or the last 10 residues (on the second plate). The average of three technical replicates per plate was used for downstream analysis.

### Data Analysis

The coefficient of determination (R^2^) was used to perform a correlation analysis between the ALS/PDC samples and the other tauopathies. Clustered heatmaps were generated using the *pheatmap* package in R, applying Euclidean distance and complete-linkage clustering to both rows and columns. A custom color scale spanning R² = 0–1 was used, and numerical R² values were displayed directly within heatmap cells to aid in comparative interpretation.

## Acknowledgments

We thank the patients’ families for donating brain tissues. This work was supported by grants from the National Institutes of Health (NIH) (1RF1AG065407-01A1) (M.I.D. and L.A.J.), the Cure Alzheimer’s Fund (M.I.D.), the Hamon Charitable Foundation (M.I.D.), AMED (JP24zf0127012, JP22jm0210097, and JP22wm0425019) (A.K.), an AARG grant from the Alzheimer’s Association, a Healey Center ALS Young Scholar Award, and a Mass General Hospital Mussallem Transformative Scholar Award in ALS Research (Y.S.), a Giovanni Armenise-Harvard Foundation award and an internal start-up fund (Y.L.).

## Notes

### Competing Interest Statement

The authors have declared no competing interest.

### Summary of Updates

Main text updated; Figures revised; Supplemental files added

