## Supplemental Figures for "Guam amyotrophic lateral sclerosis/parkinsonism-dementia complex (ALS/PDC) features CTE-like tau seeds in brain and spinal cord"

**Figure S1**

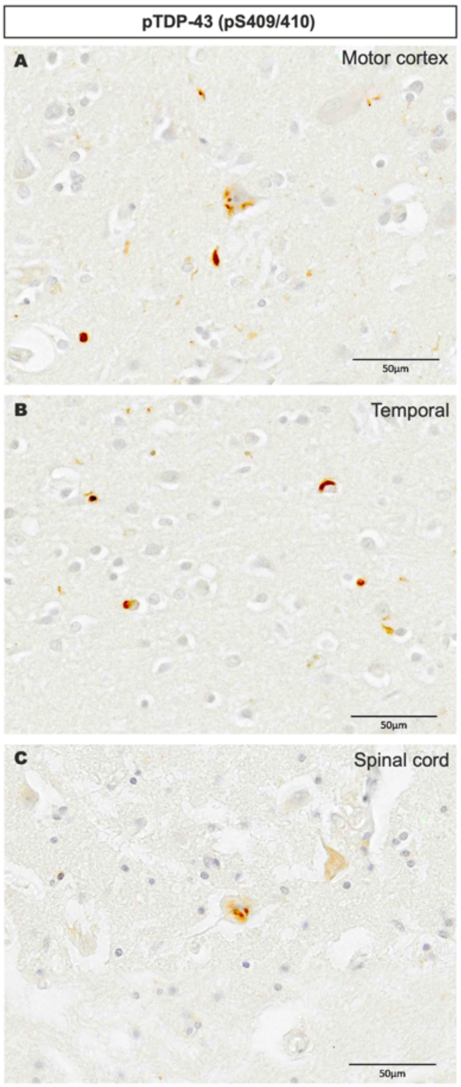


**Figure S1 | Phospho-TDP-43 aggregates in Guam ALS/PDC nervous tissues.**
Phosphorylated TDP-43 (pTDP-43) cytoplasmic inclusions can be frequently found in brain (**A**, **B**) and anterior horn of the spinal cord (**C**) (pTDP-43 pS409/410; CosmoBio, Tokyo, Japan). Tissue sections correspond to subject #3 (A, B) and subject #5 (C) from (1). We thank Dr. Tomoyo Hashimoto (University Hospital, University of Occupational and Environmental Health, Japan) for assistance in obtaining neuropathological data.

**Figure S2**

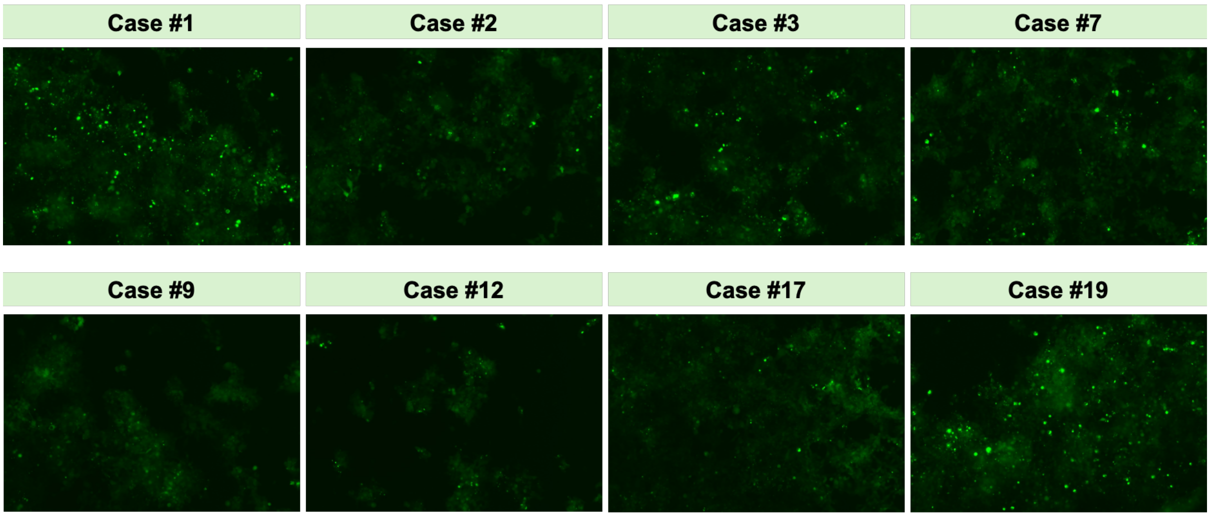


**Figure S2 | Seeding of Guam ALS/PDC tissue homogenates in 4R/4R tau biosensors.** Example images (fluorescence microscopy) of seeding results for Guam tissue homogenates in 4R/4R tau biosensor cells.

### Figure S3

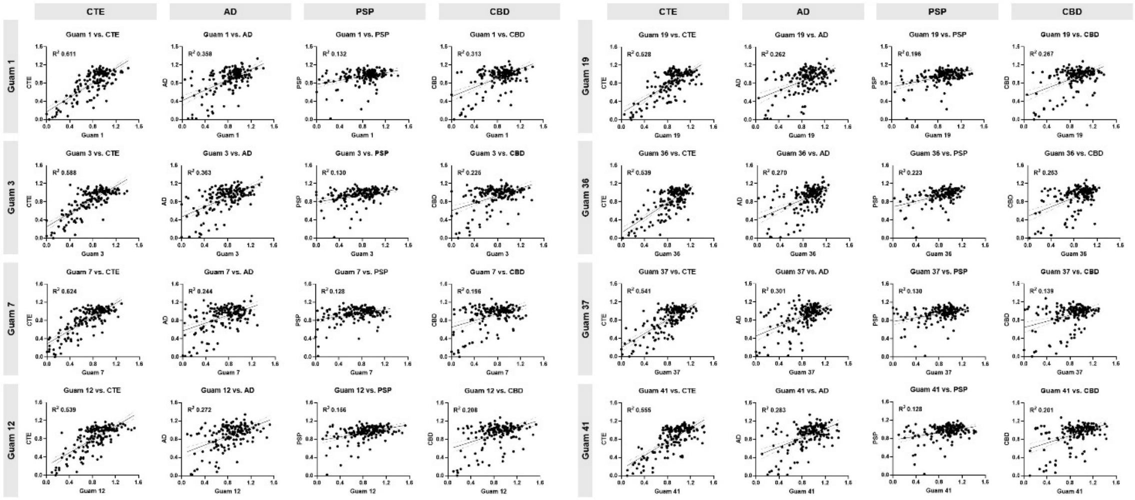


**Figure S3 | Incorporation profile comparison for ALS/PDC cases.** The incorporation profile signature for the ALS/PDC samples was compared with the average incorporation profile of CTE (n=2), AD (n=8), PSP (n=5) and CBD (n=8). The incorporation profile for these samples correlated most closely with the profile in CTE ($R^2_{Avg}$ 0.566 +/- SD 0.037), in comparison to AD ($R^2_{Avg}$ 0.294 +/- SD 0.044), PSP ($R^2_{Avg}$ 0.165 +/- SD 0.039), and CBD ($R^2_{Avg}$ 0.226 +/- SD 0.054).

**Figure S4**

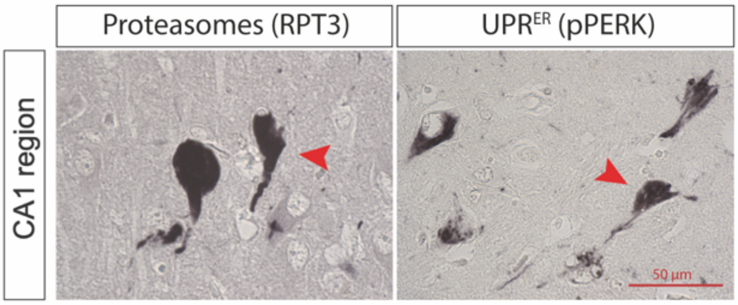


**Figure S4 | Proteasome- and pPERK-positive aggregates in Guam ALS/PDC tissues.**
ALS/PDC aggregates are immunoreactive for proteasomes (RPT3, Biomol) (2) and phosphorylated protein kinase-like endoplasmic reticulum kinase (pPERK, Santa Cruz), a marker for activation of the unfolded protein response of the endoplasmic reticulum ($UPR^{ER}$) (3). This suggests abnormalities related to intracellular proteolytic machinery and the integrated stress response pathway. pPERK-positive granules likely represent granulovacuolar bodies (4-6). Representative images of the Cornu Ammonis 1 (CA1) region of hippocampus are shown. Tissue sections correspond to subject #4 (RPT3) and subject #1 (pPERK) from (1).

**Figure S5**

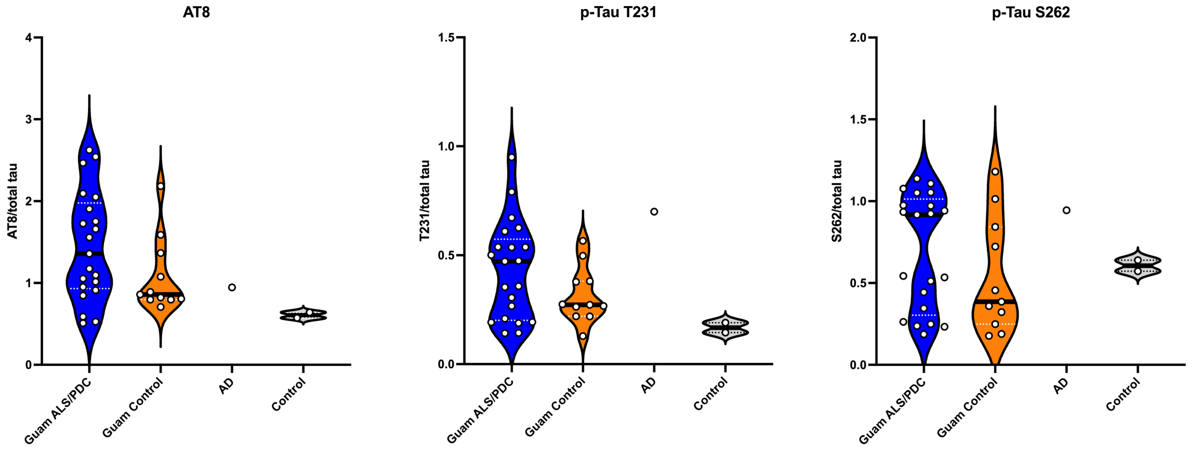


**Figure S5 | Phospho-tau modifications in Guam ALS/PDC.**

Phosphorylated tau (p-tau) modifications in Guam ALS/PDC and Guam control brain and spinal cord lysates. Dot blots were performed for p-tau (AT8/pS202+pT205, pT231, pS262) over total tau. AD: Alzheimer's disease.

### Supplementary methods

*Dot blots*

Supernatant from nervous tissue homogenates was collected for analysis. Dot blotting was performed using a vacuum-assisted 96-well dot blot apparatus and pre-wetted nitrocellulose membrane. After assembly and equilibration with Tris-buffered saline (TBS), 1 µg of each sample was diluted in 100 µl TBS and applied to the membrane. Samples were allowed to pass through by gravity filtration for 30 min, followed by vacuum filtration to complete transfer. Wells without samples were loaded with TBS alone. Membranes were then removed, blocked in blocking buffer (927-60001, LI-COR) for 1 hour at room temperature, and incubated overnight at 4°C with primary antibodies against total tau (D1M9X at 1:1000, 46687 from Cell Signaling Technology, or Tau-5 at 1:1000, ab80579 from Abcam) or phosphorylated tau (AT8 at 1:500, MN1020 from Thermofisher; phospho-tau T231 at 1:1000, 71429 from Cell Signaling Technology; or phospho-tau S262 at 1:1000, ab131354 from Abcam). Next, membranes were washed three times with Tween-20/TBS (TBS-T), incubated with secondary antibodies (anti-mouse IRDye 680RD [926-68072], anti-rabbit IRDye 680LT [926-68023], anti-mouse IRDye 800CW [926-32212], or anti-rabbit IRDye 800CW [926-32213] from LI-COR), diluted 1:20,000 for 1 hour at room temperature, washed again with TBS-T three times and rinsed with fresh TBS, before imaging on a LI-COR Odyssey imaging system. Dot blot signals were quantified by background subtraction using LI-COR Image Studio and ImageJ/Fiji, and ratios of modified tau and total tau were calculated. Data was plotted using GraphPad Prism 10.
